# Reversible disruption of specific transcription factor-DNA interactions using CRISPR/Cas9

**DOI:** 10.1101/282681

**Authors:** Ali Shariati, Antonia Dominguez, Marius Wernig, Lei S. Qi, Jan M. Skotheim

**Affiliations:** Department of Biology, Stanford University, Stanford, CA, 94305, USA; Department of Bioengineering, Stanford University, Stanford, CA 94305, USA; Department of Pathology, Stem Cell Institute, Stanford, CA 94305, USA; Department of Chemical and Systems Biology, Stanford University, Stanford, CA 94305, USA; Stanford ChEM-H, Stanford University, Stanford, CA 94305, USA

## Abstract

The control of gene expression by transcription factor binding sites frequently determines phenotype. However, it is difficult to determine the function of single transcription factor binding sites within larger transcription networks. Here, we use deactivated Cas9 (dCas9) to disrupt binding to specific sites. Since CRISPR guide RNAs are longer than transcription factor binding sites, flanking sequence can be used to target specific sites. Targeting dCas9 to an Oct4 site in the *Nanog* promoter displaced Oct4 from this binding site, reduced *Nanog* expression, and slowed division. In contrast, disrupting the Oct4 binding site adjacent to *Pax6* upregulated its transcription. Thus, we can distinguish between activating and repressing Oct4 binding sites and multiple guide RNA expression allows simultaneous inhibition of multiple binding sites and conditionally-destabilized dCas9 allows rapid reversibility. The method to disrupt specific transcription factor-DNA interactions, termed CRISPRd, is a novel high-throughput approach to systematically interrogate transcription factor binding site function within complex regulatory networks.

## Introduction

Binding of transcription factors (TFs) to specific regulatory sequences control when and where target genes are expressed. Recent technological advances have extensively mapped TF binding sites across the genome. Important as they are, these methods only provide correlative information and the function of specific binding sites remains largely unknown. It is difficult to determine the function of specific binding sites by changing TF concentration. This is because TF concentration changes will not only affect the gene of interest, but also affect hundreds of additional genes regulated by the same TF that could also contribute to the phenotype of interest (Fig. 1A).

The difficulty of determining the function of specific regulatory sites on the genome may be alleviated using CRISPR-Cas9, which can be easily programmed to target specific genomic sequences (Montalbano et al., 2017). Most commonly, CRISPR-Cas9 is used to target specific binding sites by introducing indel mutations. However, Cas9-induced mutations are random, irreversible, and lack temporal control so that lethal mutations cannot be studied (Canver et al., 2015; Diao et al., 2017; 2016; Gasperini et al., 2017; Rajagopal et al., 2016; Sanjana et al., 2016).

One possibility to alleviate the drawbacks of using catalytically active Cas9 to target specific TF-DNA binding is to use a catalytically inactive Cas9 (dCas9). Inactive dCas9 has previously been used to downregulate transcription, but not to interrogate the function of a specific TF binding site. For example, in bacteria a method known as CRISPRi is based on recruiting dCas9 to interfere with transcription machinery directly (Qi et al., 2013). In mammalian cells, a variation of this approach is based on targeting dCas9 fused with chromatin modifying enzymes that inhibit gene expression, such as the histone deacetylase complex KRAB (Dominguez et al., 2016; Klann et al., 2017; Korkmaz et al., 2016; Thakore et al., 2015; Xie et al., 2017). While such chromatin modification approaches are suitable for inhibiting gene expression, they cannot be used to determine the function of specific TF-DNA binding sites (Fig. 1B) (Groner et al., 2010; Thakore et al., 2015). More specifically, if dCas9-KRAB is targeted to successfully inhibit expression of a gene, this does not give insight into how any TF-DNA binding sites in the gene promoter regulate expression (Table 1) (Dominguez et al., 2016; Qi et al., 2013).

**Table 1.** comparison of CRISPRd with previously used CRISPR technologies in mammalian cells

| Method | Application | Mechanism | Device |
| --- | --- | --- | --- |
| CRISPRi | Repression of Transcription | Transcription block or chromatin modifications | dCas9-KRAB |
| CRISPR/Cas9 | Mutation in the regulatory sequences | Double strand break and repair | Cas9 |
| CRISPRd | Disturbing TF-DNA interaction | Competition for specific TF target site and steric hindrance | dCas9 |

## Design

To better determine the function of specific transcription factor binding sites, we developed a technique we termed CRISPRd. This method disrupts specific TF-DNA interactions by using dCas9 to sterically hinder TF binding (Fig. 1A). In contrast to CRISPRi approaches that only inhibit target gene expression without giving any information about the function of specific TF-DNA interactions, interrupting a TF-DNA interaction using CRISPRd determines whether that interaction is activating, repressing, or has no function (Fig. 1B-C; Table 1). CRISPRd works because CRISPR guide RNAs are longer than most transcription factor binding sites so that flanking sequence can be used to disrupt specific sites in the genome without affecting other sites (Fig. 2A). We also hypothesized that dCas9 can outcompete transcription factors for binding to a specific site because the dCas9 dissociation rate is significantly slower (hours) (Jones et al., 2017) than the dissociation rate of typical transcription factors (seconds) (Chen et al., 2014).

As a case study for our technology, we decided to target the Oct4 binding site 137 to 151 bp upstream of the *Nanog* transcription start site. We chose this site because of its putative role in the transcriptional positive feedback loop through which the core pluripotency transcription factors, Oct4, Nanog and Sox2, directly promote each other’s transcription to maintain pluripotency in embryonic stem cells (ESCs) (Fig. 2B). This positive feedback loop may work through the direct binding of Oct4 and Nanog to each other’s promoter (Loh et al., 2006; Young, 2011). This hypothesis is supported by reporter assays using plasmids containing 406bp of the *Nanog* promoter with and without the Oct4 binding site (Rodda et al., 2005), and by the fact that Oct4 binds to this site in ESCs (Supplementary Fig. 1A) (Kim et al., 2008; Loh et al., 2006). However, standard knockdown or depletion experiments do not directly test the positive feedback hypothesis because these three TFs occupy about 40000 sites in the mouse genome forming a highly complex transcription network (Sorrells and Johnson, 2015). Thus, that Oct4 depletion results in decreased *Nanog* expression could be an indirect consequence of the down-regulation of another Oct4 target gene.

## Results and Discussion

### CRISPRd specifically disrupts an Oct4 binding site upstream of Nanog

To test the hypothesis that Oct4 directly regulates *Nanog* expression. We designed single guide RNAs (sgRNAs) that target dCas9 to the Oct4 binding site (Fig. 2C-D). We used lentiviral infection to express these sgRNAs in mouse ESCs expressing a Dox-inducible dCas9-mCherry fusion protein (Fig. 2E) and in which one allele of *Nanog* was tagged with Venus at the endogenous locus (*Nanog-Venus/dCas9-mCherry mESC*). We designed two sgRNAs, Oct4-Site Nanog-1 and Oct4-Site Nanog-2, that bind to the 14 bp Oct4 binding site and an additional 8 or 10 bp flanking sequence respectively. We note that the flanking sequence confers specificity to the Oct4 binding site in the *Nanog* promoter over other Oct4 binding sites in the genome (Fig. 2C). We also included 4 other control sgRNAs (Control1-4) that do not overlap with the targeted Oct4 binding site (Fig. 2D).

Expression of the sgRNAs targeting the Oct4 site decreased Nanog-Venus protein level as measured by flow cytometry (Fig. 3A). Similarly, the mRNA level of Nanog decreased by more than 50% when dCas9 was targeted to the Oct4 binding site (Fig. 3B). In contrast, expressing control sgRNAs, Control1-4, targeting dCas9 either upstream or downstream of the target site did not have this effect (Fig. 2D, 3B). Both wild type Nanog and Nanog-Venus proteins show reduced expression, which implies that dCas9 can displace Oct4 from its binding site on both Nanog alleles (Fig. 3C). To directly test if dCas9 recruitment sterically hinders the binding of Oct4, we used chromatin immunoprecipitation (ChIP) to measure the binding of Oct4 and dCas9 in cell lines expressing targeting or control sgRNAs. Expression of targeting sgRNAs significantly reduced the binding of Oct4 to its target site, while increasing the binding of dCas9 to this same site. This was not the case for control sgRNAs (Fig. 3D-E and Supplementary Figure 1B).

We next sought to compare the effect of using CRISPRd to block Oct4 access to its binding site in the *Nanog* promoter with the effect of deleting this site using active Cas9. To do this, we deleted the Oct4 binding site upstream of the *Nanog-Venus* allele using active Cas9. Deletion of this site reduced Nanog-Venus expression by ~2-fold, which was comparable to the effect of inhibiting Oct4 binding to the site with CRISPRd (Fig. 3F; Supplementary Fig. 1C). These findings suggest that this specific Oct4 binding site can contribute to about 50% of *Nanog*mRNA transcription and CRISPRd can efficiently counteract the effect of this site on *Nanog*expression by blocking Oct4 binding.

To test the functional outcome of interfering with Oct4 binding to the *Nanog* promoter, we used time-lapse imaging to measure cell cycle progression in individual mESCs. Disrupting Oct4 binding to the *Nanog* promoter elongated the cell cycle (median cell cycle duration increased from 12h-13h to 16-17h; Fig. 3G, Supplementary Videos 1 and 2). This is consistent with the previously reported slow growth phenotype of Nanog deficient ESCs (Chambers et al., 2003; Mitsui et al., 2003). Thus, our experiment strongly supports the hypothesis that Oct4 directly promotes *Nanog* expression in mESCs to accelerate cell proliferation. Importantly, this example demonstrates that CRISPRd can be used to sterically hinder TF binding on specific sites on the genome in mammalian cells, which can be used to determine the contribution of individual TF binding sites to specific cellular functions.

## CRISPRd can distinguish between activating and repressing binding sites

Unlike CRISPRi, which solely aims to inhibit transcription by modifying local chromatin, CRISPRd can be used to determine if a specific TF-DNA binding site promotes or inhibits transcription. Above, we used CRISPRd to show that Oct4 binding to the Nanog promoter promoted transcription. Here, we examine a case where an Oct4-DNA interaction inhibits transcription. In addition to binding to the promoters of genes involved in self-renewal, Oct4 binds to the promoters of genes regulating lineage specification, including the neuronal lineage (Loh et al., 2006). Interestingly, Oct4 also associates with transcriptional repressors in pluripotent cells (Liang et al., 2008), which suggests that Oct4 can both activate self-renewal genes and repress neuronal target genes to suppress neuronal fate in embryonic stem cells (Thomson et al., 2011). If this is the case, blocking an Oct4 repressive binding site near its neuronal targets should result in increased expression of the target gene. *Pax6* is a pan-neuronal transcription factor that is not transcribed in mouse ESC but whose promoter is bound by Oct4 (Fig. 4A) (Kim et al., 2008). Disruption of this binding site with CRISPRd resulted in a 5±1-fold increase in *Pax6* mRNA (se=0.7) and an increase in Pax6 protein (Fig. 4B-C; similar results were obtained for a second sgRNA). These findings show how CRISPRd can determine the function of a TF-DNA binding site that inhibits transcription.

## CRISPRd allows simultaneous targeting of multiple TF binding sites

The ease with which multiple sgRNAs can be expressed in cells suggested that we can extend our approach to simultaneously target multiple TF binding sites. This multiplexing can be used to interrogate complex transcription networks in which phenotypes result from several TF target genes. To extend our approach, we chose to target another Oct4 binding site that was shown to regulate the *Utf1* gene in reporter assays (Kooistra et al., 2010). *Utf1* itself is a transcription factor that regulates chromatin organization in mESCs and it shares many targets with Oct4 (Kooistra et al., 2010). Oct4 binds to its regulatory sequence downstream of the gene *Utf1*, which is consistent with the presence of an Oct4 ChIP-Seq binding peak (Fig. 5A, Supplementary Fig. 2A-B). Recruitment of dCas9 to this Oct4 site using three different sgRNAs (Oct4-Site Utf1 -1-3) reduced *Utf1* mRNA expression, reduced Oct4 binding, and increased dCas9 binding to the targeted Oct4 site (Fig. 5B-E). Next, we infected cells with lentiviral particles to express two sgRNAs from a dual guide RNA construct (Fig. 5F). Targeting dCas9 to the Oct4 sites in the regulatory sequence of both *Nanog* and *Utf1* genes reduced *Nanog* mRNA and protein levels, and reduced *Utf1* mRNA levels (Fig. 5G-H). Importantly, these sgRNAs did not result in a significant decrease in Oct4 binding to two other Oct4 binding sites with similar binding motifs but with different flanking sequence (Supplementary Fig. 1C-F). These results demonstrate that dCas9 can be used to simultaneously disrupt TF binding at multiple loci.

## Reversible disruption of TF binding using destabilized dCas9

While targeting active Cas9 to TF binding sites can also be used to ascertain their function, such mutagenic approaches are not reversible. Reversibility is important because it allows the study of essential transcription factor binding sites, and can be used to generate controllable dynamics of inhibition of a specific site at a specific time. For example, one could inhibit a site during a specific interval of development or cell cycle phase. Our approach can be modified to make it rapidly reversible. To do this, we employed a conditionally destabilizing domain (DD) that is rapidly degraded in the absence of the small molecule Shield1 (Banaszynski et al., 2006). We generated mESCs expressing dCas9 fused to a DD domain and mCherry (ddCas9) (Fig. 6A). To measure the kinetics of ddCas9 degradation, we first grew ddCas9 expressing ESCs in the presence of Shield1. Next, we titrated out Shield1 by adding excess amounts of the purified destabilized domain to the media (Miyazaki et al., 2015). Most of the ddCas9-mCherry is degraded in less than one hour as measured by immunoblotting and live-cell imaging (Fig. 6B-C, Supplementary Video 3). To determine how rapidly transcription can be altered using our ddCas9 approach, we infected the *ddCas9-mCherry* ESC line with a virus containing an sgRNA (Oct4-Site Nanog-1) that targets the Oct4 site in the *Nanog* promoter. After destabilizing ddCas9, Nanog mRNA increased more than 5-fold within one hour, which was closely followed
by an increase in Nanog protein (Fig. 6D-E). These results show that ddCas9 can be used to rapidly and reversibly interfere with transcription factor binding.

Here, we showed that dCas9 can compete with endogenous transcription factors to disrupt their binding to specific target sites. This approach can be easily multiplexed to simultaneously target multiple TF sites and the fusion of a conditionally destabilized domain to dCas9 allows rapid and reversible exogenous control of TF binding to specific sites. We expect our approach to determine the function of specific TF binding sites within complex transcription networks via systematic perturbation using a library of sgRNAs.

## Limitations

CRISPRd is limited by the presence of PAM sequences near the TF binding site of interest. This can be alleviated by Cas9 orthologs with different PAM requirements (Dominguez et al., 2016; Shalem et al., 2015). It should also be noted that because of slow search kinetic of dCas9 (Jones et al., 2017), disrupting a TF-DNA interaction is not immediate and can take hours. Protein engineering techniques to generate dCas

## Methods and Materials

### Cloning

sgRNAs were expressed using a lentiviral mouse U6 (mU6)-based expression vector that coexpressed Puro-T2A-BFP from an EF1α promoter (Fig. 2E). New sgRNA sequences were generated by PCR and introduced by InFusion cloning into the sgRNA expression vector digested with BstXI and XhoI. For multiplexing experiments, sgRNAs were expressed using a lentiviral dual sgRNA vector consisting of two sgRNA cassettes in tandem driven by the human U6 promoter and mouse U6 promoter, respectively, and a Puro-T2A-BFP cassettes (Fig. 4F). In the duel sgRNA vector, the mU6 vectors are cloned using InFusion to insert PCR products into a modified vector digested with BstXI and XhoI. The hU6 sgRNA vector was cloned by inserting PCR products with InFusion cloning into the parent vector digested with SpeI and XbaI. After sequence verification, the mU6 vector was digested with XbaI and XhoI and the mU6 sgRNA cassette was ligated into the hU6 vector digested with SpeI and SalI.

To assemble the doxycycline-inducible dCas9 construct (pSLQ1942), human codon-optimized *S. pyogenes* dCas9 was fused at the C-terminus with an HA tag and two SV40 nuclear localization signals (Fig. 2E). For visualization, mCherry was fused at the C-terminus following a P2A peptide. This cassette is driven by the TRE3G doxycycline-inducible promoter. Zeocin resistance and Tet-On 3G transactivator expression is driven by the Ef1α promoter. These cassettes were cloned into a Piggybac plasmid containing the 5’ and 3’ Piggybac homology arms. To assemble the ddCas9 plasmid (pSLQ2470), the destabilization domain that can be stabilized by Shield1 was amplified from pBMN FKBP(L106P)-YFP-HA and inserted into pSLQ1942. Then, the IRES HcRed-tandem was inserted using InFusion Cloning into KpnI digested pSLQ1942.

## Cell culture

All the ES lines were grown in ESGRO-2i medium (SF016-200, Millipore) supplemented with 100 units/ml streptomycin and 100 mg/ml penicillin on cell culture dishes coated with 0.1% gelatin (G1890, Sigma-Aldrich). The media was changed everyday and cells were passaged every 2 days using Accutase cell detachment solution (SCR005, Millipore). The HEK293T cells were grown in DMEM/F12 supplemented with 10% Fetal Bovine Serum (FBS), 100 units/ml streptomycin, and 100 mg/ml penicillin. All cell culture experiments were done at 37°C and 5% CO_2_.

## Generation of Nanog-Venus/dCas9-mCherry cell line

To generate an inducible dCas9 expressing ESC cell line, we inserted tetracycline inducible dCas9-mCherry into the genome of a Nanog-Venus-mESC line using the piggybac transposon system (Ding et al., 2005; Filipczyk et al., 2013) (Fig. 2E). Cells were transfected with a tet-on dCas9 plasmid (pSLQ1942) and PiggyBac transposase using the Turbofect transfection reagent following the manufacturer’s instructions (R0531, ThermoFisher Scientific). A clonal line was generated by manually selecting an mCherry positive colony and expanding the colony in 2i medium. Addition of 1μg/ml doxycyclin to the medium of the Nanog-Venus/dCas9-mCherry line resulted in a 65-fold increase in the dCas9-mCherry protein signal while the Nanog-Venus protein signal did not change (Supplementary Fig. 3A-B). A similar strategy was used to infect Nanog-Venus line with destabilized dCas9 vector,pSLQ2470, to generate the Nanog-Venus/ddCas9-mCherry.

## Deletion of Oct4 binding site

We used Edit-R lentiviral inducible Cas9 (Dharmacon) to generate a Nanog-Venus-mESC line that expresses Cas9 by addition of doxycyclin. Next, we infected this line with lentiviral particles encoding an sgRNA targeting slightly downstream of the Oct4 binding site upstream of *Nanog*. The cells were grown in the presence of doxycyclin (1μg/ml) for three days. Individual colonies were grown in a 96 well plate and were genotyped using the following primers: Nanog-Genotype-F (5′-CTTCTTCCATTGCTTAGACGGC-3′), Nanog-Genotype-R (5′-GGCTCAAGGCGATAGATTTAAAGGGTAG-3′). We sequenced the PCR products of the genotyping reaction and a line with a ~230 bp deletion including the Oct4 binding site upstream of Nanog was used for analysis.

## sgRNA lentiviral production

Lentiviral particles containing sgRNA expression plasmids were generated by transfecting HEK293T cells with sgRNA plasmids, and with standard packaging constructs using the Turbofect transfection reagent as previously described (Schwarz et al., 2018) (R0531, ThermoFisher Scientific). One day after transfection, the HEK293T cell media was changed from DMEM/FBS to 2i (SF016-200, Millipore). The viral particles in the 2i media were collected after 48 hours, centrifuged, and filtered (0.45μm syringe filter). The particles were then added to media of the Nanog-Venus/dCas9-mCherry ESC line. The sgRNA expressing cells were selected using puromycin. The expression of sgRNAs was also visually confirmed by microscopy of BFP expression. Supplementary Table 1 shows the sequences of all the sgRNAs used in this study.

## Oct4 binding site identification

To identify Oct4 binding sites, we used available ENCODE ChIP-seq data to find an Oct4 peak near the transcription start site (TSS) of *Nanog*. This broad Oct4 peak is between 500 and 33 bp upstream of the *Nanog* TSS (Fig. 2D). To find the exact location of the Oct4 binding site, we searched for transcription factor motifs using the JASPAR database (Sandelin, 2004). This identified a consensus binding site between 137 and 151 bp upstream of the *Nanog* TSS. A similar strategy was used to identify the Oct4 binding site located 1825 bp downstream of the *Utf1* TSS (Fig. 4A). sgRNAs were designed based on available PAM sites (NGG) near the Oct4 binding sites. To obtain the vertebrate TF binding site length, we used the TFBSTools bioconductor package to access the vertebrate TF binding site position frequency matrices of the JASPAR2018 library dataset (Khan et al., 2018; Tan and Lenhard, 2016). The number of columns per matrix was used to obtain the distribution of the vertebrate TF binding site length.

## Chromatin immunoprecipitation-qPCR (ChIP-qPCR)

ChIP-qPCR was performed using a SimpleChIP Enzymatic Chromatin IP kit following the manufacturer’s protocol (#9002, Cell Signaling Technology). Briefly, up to 4 X 10^7^ cells were fixed using 4% paraformaldehyde, and chromatin was prepared and fragmented using Micrococcal nuclease (Mnase). Duration and enzyme concentration was optimized to obtain chromatin fragments between 150 and 900 bp. Fragmented chromatin was incubated overnight at 4°C with antibodies against HA or Oct4 to pull down dCas9-HA or Oct4 on ChIP grade agarose beads. Beads were washed several times, DNA-Protein cross-linking was reversed, and DNA was purified on a column. The purified DNA was used to quantify the binding of Oct4 or HA-dCas9 relative to input using quantitative PCR (qPCR). A list of primers used for ChIP-qPCR analysis is provided in Supplementary Table 2. qPCR was performed with two technical replicates and three or four biological replicates. All the reported enrichment values are normalized to the experiment done on a line expressing HA-dCas9, but no sgRNA. The enrichment was calculated by subtracting the Ct value from qPCR of the pull-down input from the Ct value from qPCR of the of chromatin input (ΔCt). The ΔCt for each sgRNA was subtracted from the ΔCt from the dCas9 only line (ΔΔCt) and relative enrichment was calculated as 2^ΔΔCt^. A goat polyclonal anti-Oct4 antibody (N19, sc-8628, Santa Cruz Biotechnology) was used to pull-down Oct4, and a rabbit polyclonal anti-HA tag (ab9110, Abcam) was used to pull down dCas9 tagged with HA and mCherry.

## Quantitative RT-PCR

Total RNA was harvested from cells using a PARIS RNA isolation kit 3 days after inducing the expression of dCas9 (AM 1921, ThermoFisher Scientific) using 1 μg/ml of doxycycline. DNA contamination was removed by treating the isolated RNA with DNAase using a TurboDNA free kit (AM 1907, ThermoFisher Scientific). Quantitative RT-PCR was performed using an iTaq Universal SYBR green one-step kit and an iq-5 Bio-Rad instrument. The primer sequence is shown in Supplementary Table 2. qRT-PCR experiments were performed with two technical replicates and at least 3 biological replicates. mRNA fold change was calculated by subtracting the ΔCt (Ct of tested gene minus Ct for Actin) in samples from lines expressing dCas9 and different sgRNAs from the ΔCt of samples from the control line only expressing dCas9, but no sgRNA (ΔΔCt). The fold change was then calculated as 2^-ΔΔCt^.

## Time-lapse microscopy

For time lapse imaging, cells were plated on 35 cm glass bottom dishes (MatTek) coated with laminin (LN-521^TM^ STEM CELL MATRIX). Imaging experiments were performed 3 days after the induction of dCas9 in a chamber at 37°C perfused with 5% CO_2_ ^25^. Images were taken every 30 minutes for cell cycle measurements and every 20 minutes for ddCas9 degradation measurements at up to 3 positions per dish for 3 dishes using a Zeiss AxioVert 200M microscope with an automated stage, and an EC Plan-Neofluar 5x/0.16NA Ph1 objective or an A-plan 10x/0.25NA Ph1 objective. Cell cycle duration was calculated by manually tracking cells from when they are born until when they complete mitosis. For ddCas9-mCherry degradation analysis, an excess amount of purified destabilized domain was added to the medium to titrate Shield1 and then the mCherry signal was measured. Cells were manually segmented to calculate the total amount of mCherry within each individual cell. Background signal from the area adjacent to the cell was measured and subtracted from the mCherry signal. The signal for each cell was then normalized to its value in the first frame of the movie.

## Immunoblot

Cells were lysed using an RIPA buffer supplemented with protease and phosphatase inhibitors. Proteins were separated on a 8% SDS-PAGE gel and were transferred to a Nitrocelluluse membrane using an iBlot (IB21001, ThermoFisher Scientific). Membranes were incubated overnight at 4°C using the following primary antibodies: polyclonal rabbit anti-Nanog antibody (A300-397A, Bethyl Laboratories Inc.), polyclonal goat anti-Oct3/4 antibody (N19, sc-8628, Santa Cruz Biotechnology), and a rabbit polyclonal anti-HA (ab9110, abcam) to detect dCas9, Mouse Alpha-Tubulin(Sigma, T9026), Monoclonal Mouse Gapdh (MA5-15738, Pierce). The primary antibodies were detected using fluorescently labeled secondary antibodies (LI-COR) and were visualized using Licor Odyssey CLx.

## Immunofluorescence staining

Cells were plated on 35 cm glass bottom dishes (MatTek) coated with laminin. Cells were grown in 2i media supplemented with doxycyclin (1μg/ml) to induce dCas9 expression. The cells were washed with PBS two times and then fixed using 4% paraformaldehyde for 10 minutes. Cells were washed in PBS and were incubated in 1.5% Bovine Serum Albumin (BSA) solution for 1h. Cells were stained with a chicken Pax6 antibody (Developmental Studies Hybridoma Bank) for 2 hours at room temperature. Alexa 633 secondary antibody (Invitrogen) was used to visualize the Pax6 signal and DAPI was used to visualize the nuclei. The intensity of Pax6 signal for each cell was measured by segmenting the nuclei and subtracting the background from an area adjacent to the cell.

## Flow cytometry

Cells were grown in the presence of doxycyclin (1μg/ml) for 3 days before exposure to an Accutase cell detachment solution. Cells were pelleted at 1000 rpm. The pellet was then resuspended in PBS and filtered using a 40μm Cell Strainer (Cornings, #352340). The flow cytometry measurements using a 488 nm Blue laser to detect Nanog-Venus were performed using a FACScan analyzer or a FACS ARIA at the Stanford University FACS facility. The flow cytometry measurements were repeated two times and for each experiment the Nanog-Venus amount of different sgRNAs was normalized to the median amount of dCas9 only expressing cells. For dual guide experiments cells were treated with doxycyclin (1μg/ml) for 5 days before flow cytometry analysis and the Nanog-Venus amounts were not normalized.

## Acknowledgements

The authors would like to thank members of the Skotheim, Qi and Wernig labs for providing feedback and reagents, and to thank Ben Topacio, Devon Brown and Dr. Evgeny Zatulovskiy for their help with flow cytometry analysis and cell culture. Flow cytometry analysis for this project was done on instruments in the Stanford Shared FACS Facility. We also thank Dr. Airlia Thompson for providing the purified DD protein, and Dr. Tom Wandless for providing the pBMN FKBP(L106P)-YFP-HA IRES HcRed-tandem plasmid. This work was supported by the NIGMS/NIH through an NRSA Award F32GM123576 (AS), ALS Association Milton Safenowitz Fellowship, Burroughs Wellcome Fund Postdoctoral Enrichment Program (AAD) and R01
GM092925 (JMS), and by Stanford University through a Bio-X Seed Grant (AS, AD, JMS, MW, LSQ).

## Author Contribution

All authors participated in experimental design. AS performed and analyzed ChIP-qPCR, qRT-PCR, microscopy, Immunoblot and flow cytometry experiments. MW helped set up the mESCs culture. AAD designed and cloned all the sgRNA, dCas9 and conditionally stable dCas9 vectors. JMS and AS wrote the paper.

**Figure 1:**
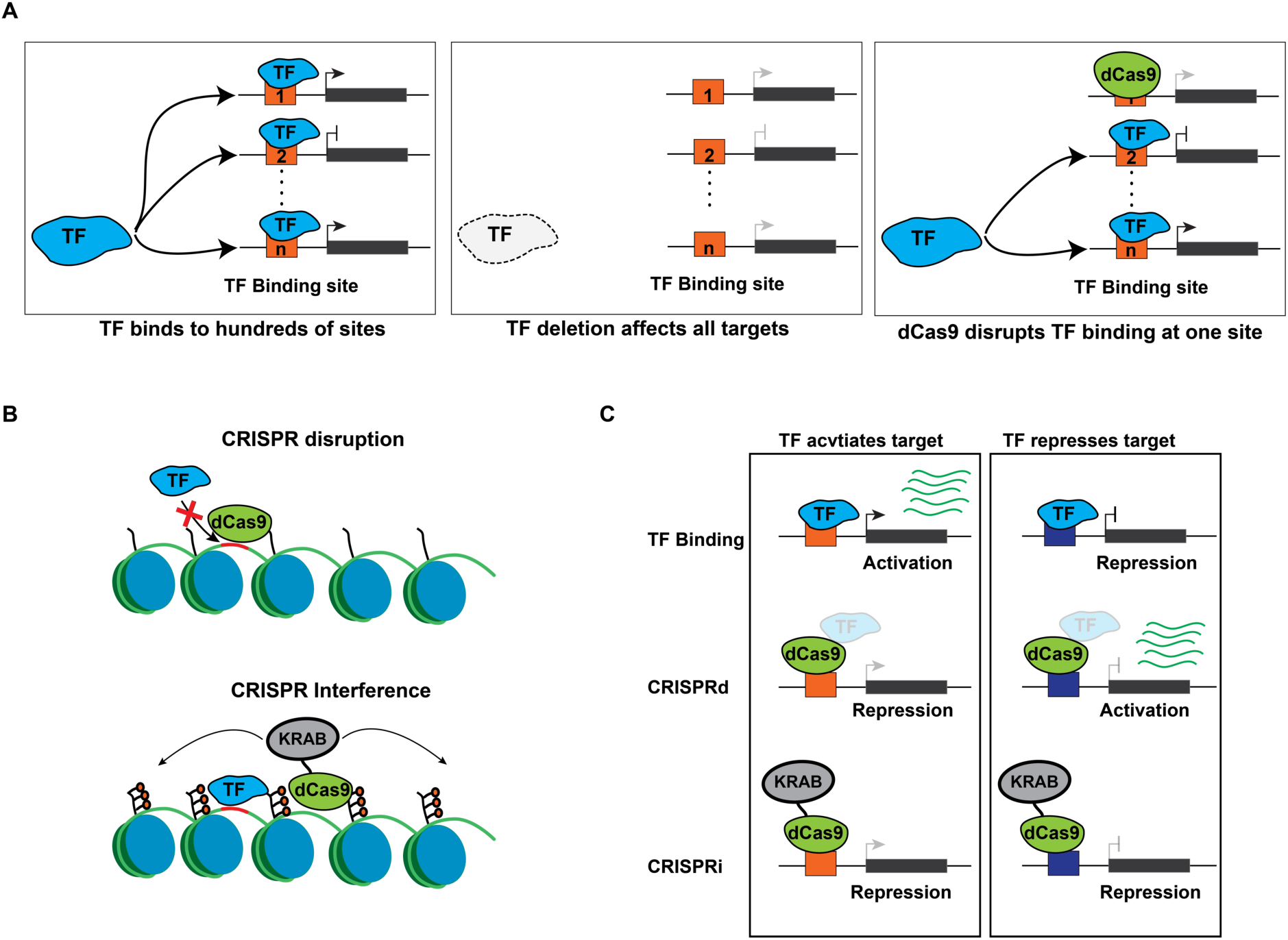
Experimental design and comparison with existing methods. A. Schematic showing a TF binding hundreds of sites across the genome (right panel). Function of individual binding sites cannot be determined by deleting the TF as this will not only affect the binding site of interest but also hundreds of other targets (middle panel). CRISPRd targets dCas9 to sterically inhibit transcription factor binding at a specific site (left panel).
B. Comparison of CRISPRi approaches, which aim to downregulate gene expression by inducing chromatin modifications, and CRISPRd, which aims to disrupt a specific TF-DNA interaction.
C. Schematic showing the expected effects on gene expression using CRISPRd and CRISPRi. A TF binding site interaction can activate or repress target genes (top panel). CRISPRd can distinguish activating from repressing functions (middle panel). CRISPRi represses the expression of the targeted locus without distinguishing between activating or repressing TF-DNA binding sites (lower panel).

**Figure 2:**
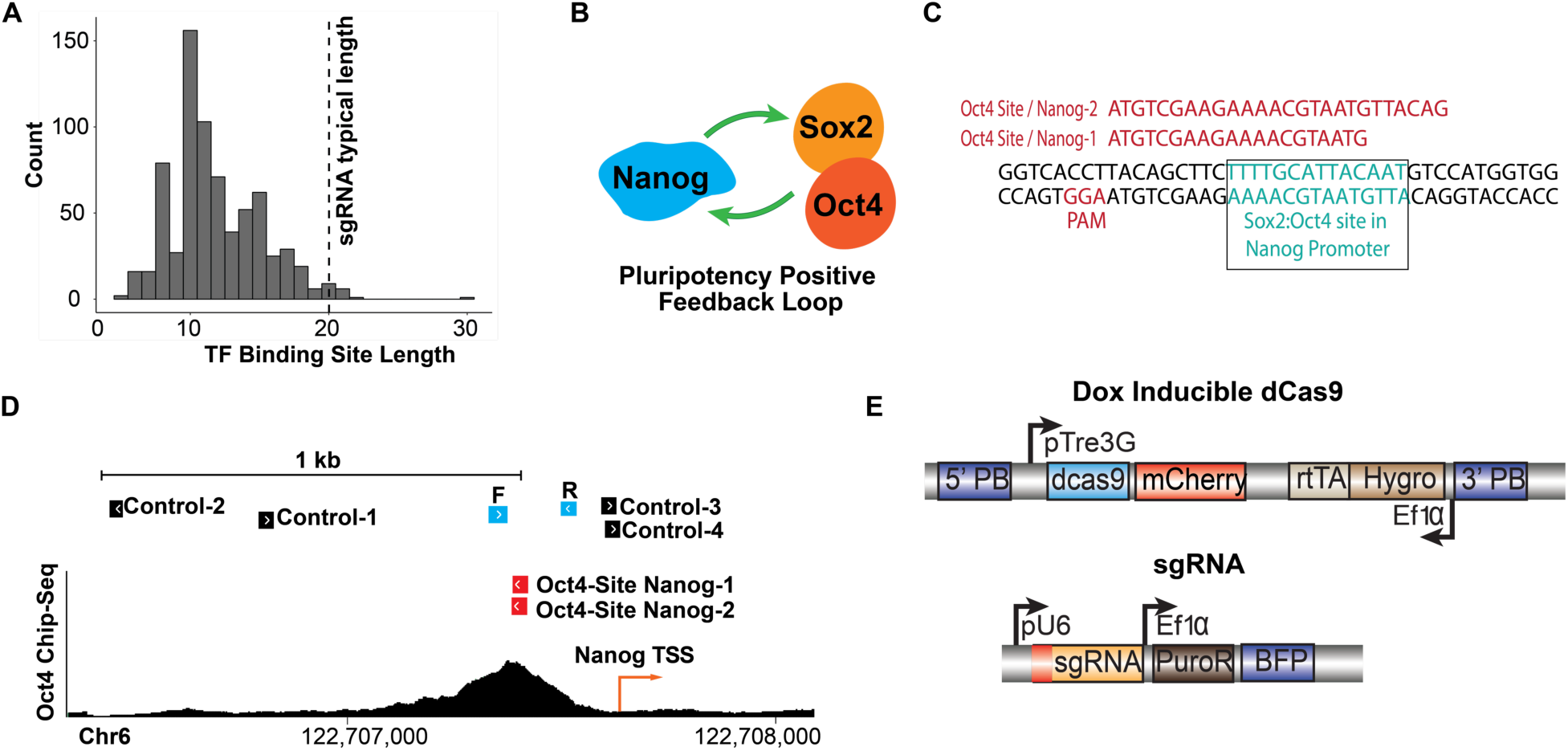
sgRNA design for CRISRPd to target the pluripotency positive feedback loop. A. Distribution of transcription factor binding site length in vertebrate genomes. The vertical line shows the typical size of an sgRNA. Typical sgRNAs are longer that most TF binding sites. The flanking sequence can be used to design specific sgRNAs against individual binding sites.
B. Schematic of the transcriptional positive feedback loop that maintains pluripotency in mouse embryonic stem cells.
C. The Oct4 binding motif upstream of *Nanog* is shown in green and the targeting sgRNAs and PAM sequences are shown in red.
D. ChIP-seq data for Oct4 show the chromosomal position of an Oct4 peak upstream of *Nanog*. Also shown are the position of targeting sgRNAs (red), Control sgRNAs (black), and qPCR primers (blue).
E. The doxycyclin-inducible vector contains dCas9 under the control of a *TRE3G* promoter and another cassette with an *EF1α* promoter driving hygromycin resistance and an rtTA transactivator. The sgRNA vector contains an sgRNA cassette with customizable guide sequence expressed from the U6 promoter and an expression cassette containing an *EF1α* promoter driving expression of a puromycin-resistance gene and BFP.

**Figure 3:**
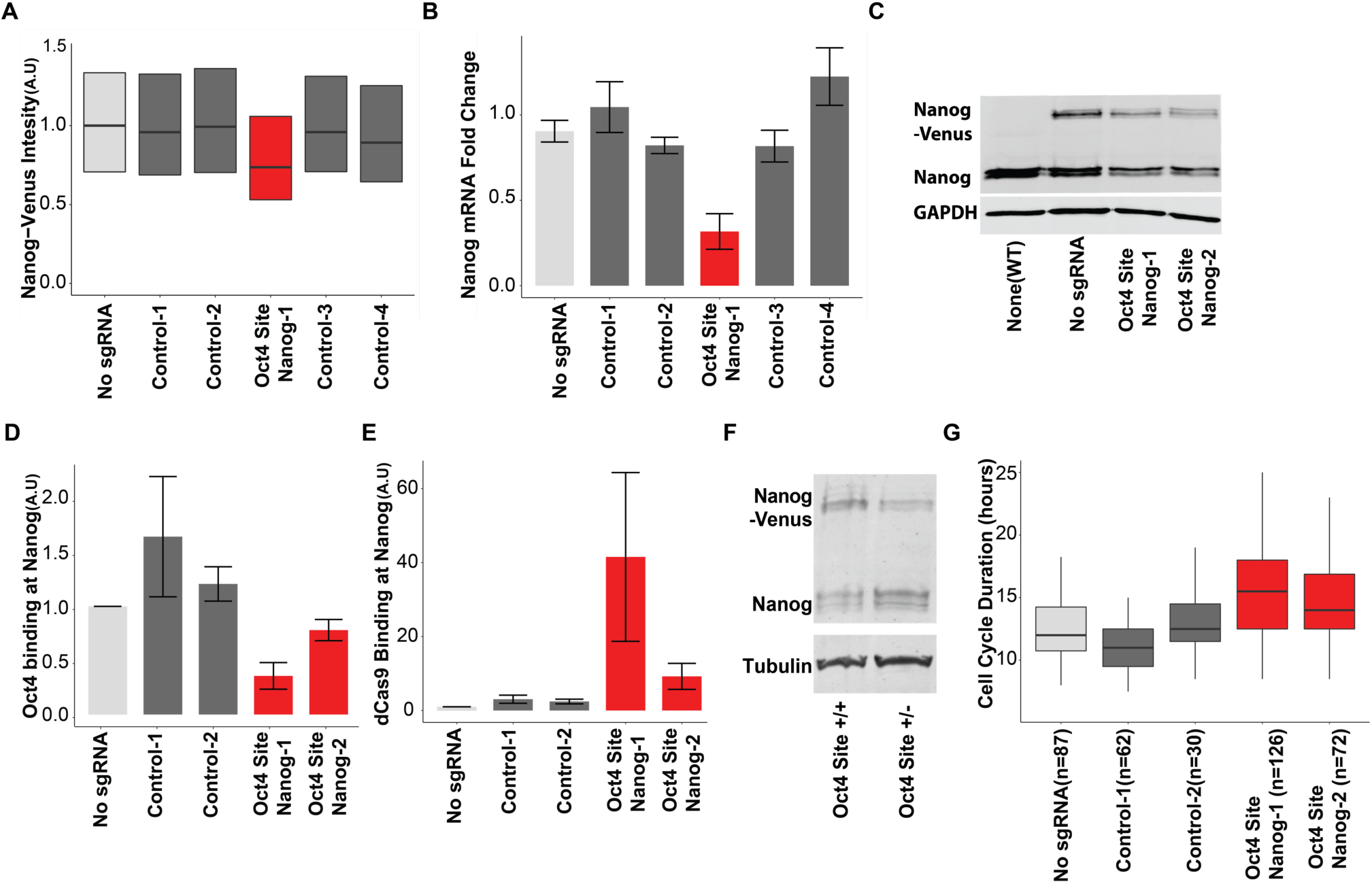
CRISPRd disrupts Oct4 binding in the *Nanog* promoter. A. Flow cytometry of Nanog-Venus shows that targeting an upstream Oct4 binding site results in Nanog mRNA reduction. Control sgRNAs targeting dCas9 upstream or downstream of the target site has no effect.
B. qRT-PCR measurement of *Nanog* mRNA shows a decrease when the targeting sgRNA is expressed, but not when the control sgRNAs are expressed.
C. Immunoblot of Nanog shows both *Nanog* alleles decrease their expression.
D. Oct4 ChIP-qPCR shows that Oct4 binding upstream of Nanog is reduced by targeting sgRNAs.
E. dCas9 ChIP-qPCR measurements for cells expressing dCas9 and the indicated sgRNA9.
F. Immunoblot of Nanog protein in wild type (+/+) and heterozygous (+/−) Oct4 binding site deletion mESCs.
G. Distributions of cell cycle durations for cells expressing dCas9 and the indicated sgRNA. For all panels, the targeting sgRNAs are shown in red, the control sgRNAs are shown in dark gray, and the no sgRNA control is shown in light gray. Bottom and upper lines of box plots show the first and third interquartile range and the middle line shows the median. Bar plots show mean and associated standard error.

**Figure 4:**
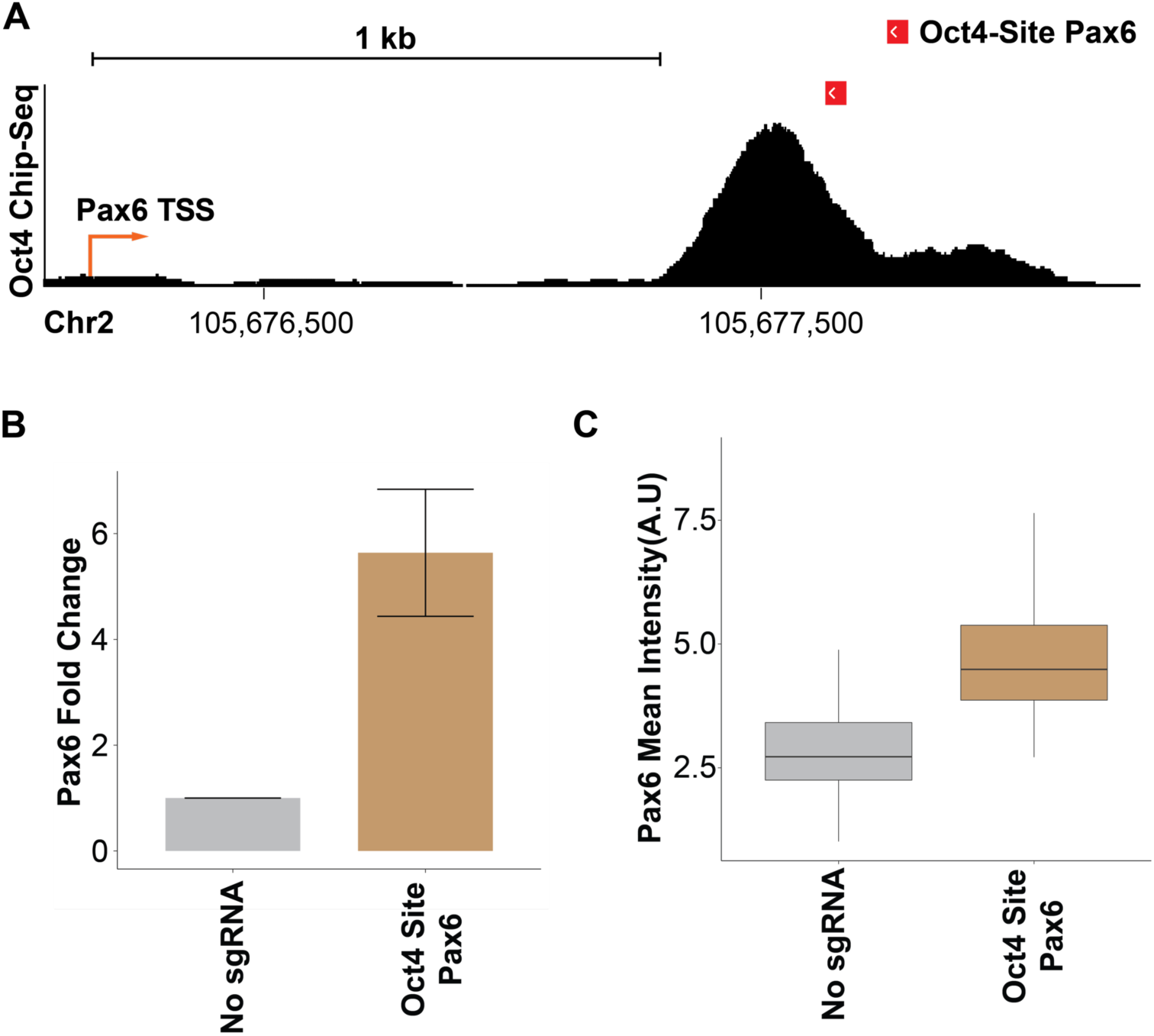
CRISPRd disrupts Oct4 inhibition of *Pax6* expression. ChIP-seq data for Oct4 show the chromosomal position of an Oct4 peak in the regulatory region of *Pax6* gene. Also shown is the position of the targeting sgRNA (red). B. qRT-PCR of *Pax6* in cells expressing the sgRNA targeting the nearby Oct4 site. C. Immunofluorescence staining of Pax6 protein. Bottom and upper lines of box plots show the first and third interquartile range and the middle line shows the median. Bar plots show mean and associated standard error.

**Figure 5:**
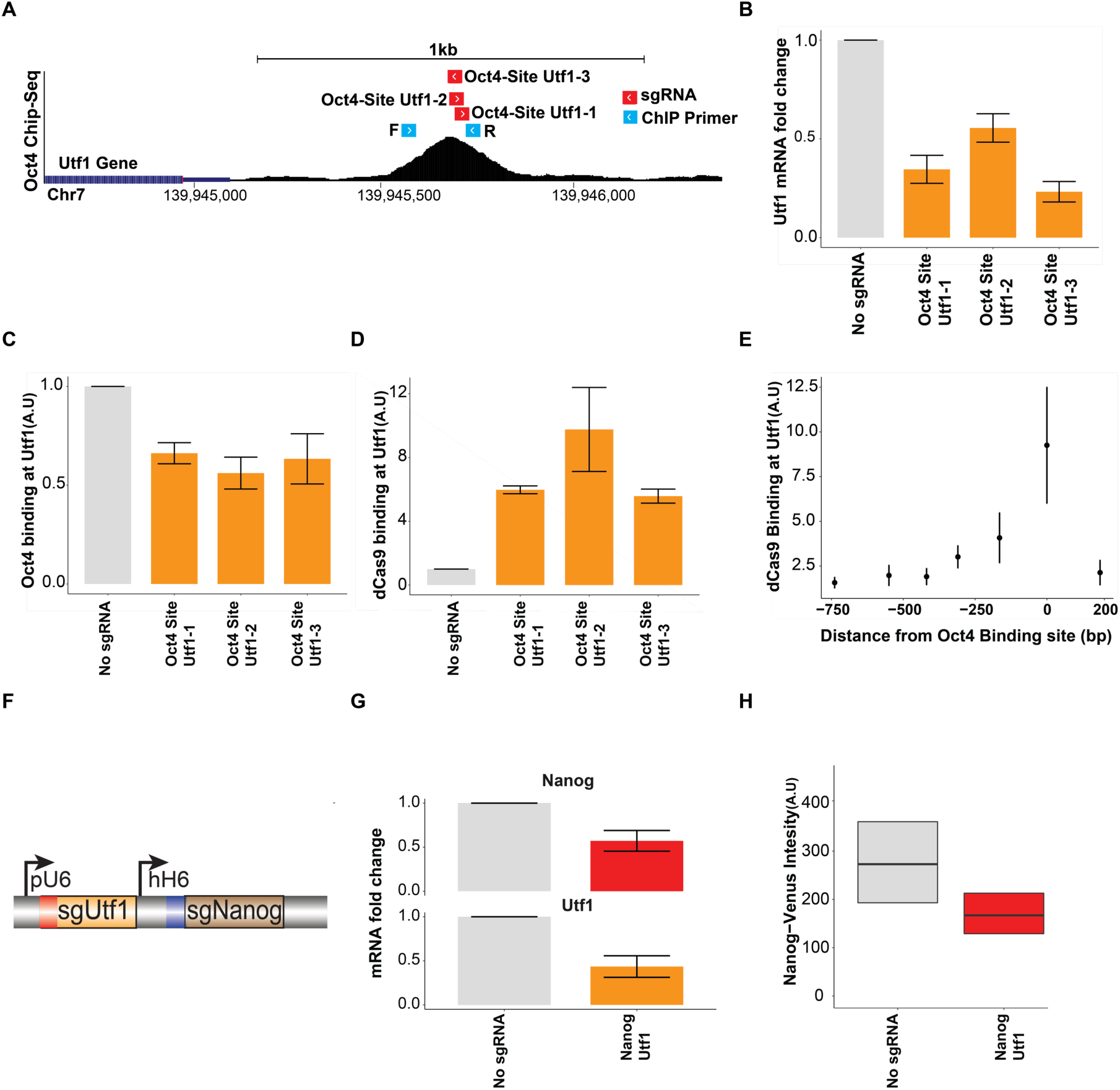
Multiplexing CRISPRd. A. Oct4 ChIP-seq data near *Utf1*. Also shown are sgRNA, and qPCR primer sequences.
B. *Utf1* mRNA qRT-PCR measurement,
C. Oct4 ChIP-qPCR, and,
D. dCas9 ChIP-qPCR for cells expressing dCas9 and the indicated sgRNAs.
E. dCas9 ChIP-qPCR measurement using overlapping primers near the Oct4 binding site downstream of the *Utf1* gene.
F. Schematic of the construct used to express two sgRNAs.
G. *Nanog* and *Utf1* mRNA qRT-PCR measurement in cells expressing either dCas9 alone or with the sgRNAs targeting the indicated Oct4 sites near both genes (Nanog Utf1).
H. Flow cytometry measurement of Nanog-Venus protein in control and Nanog Utf1 sgRNA expressing cells. Bottom and upper lines of box plots show the first and third interquartile range and the middle line shows the median. Bar plots show mean and associated standard error.

**Figure 6:**
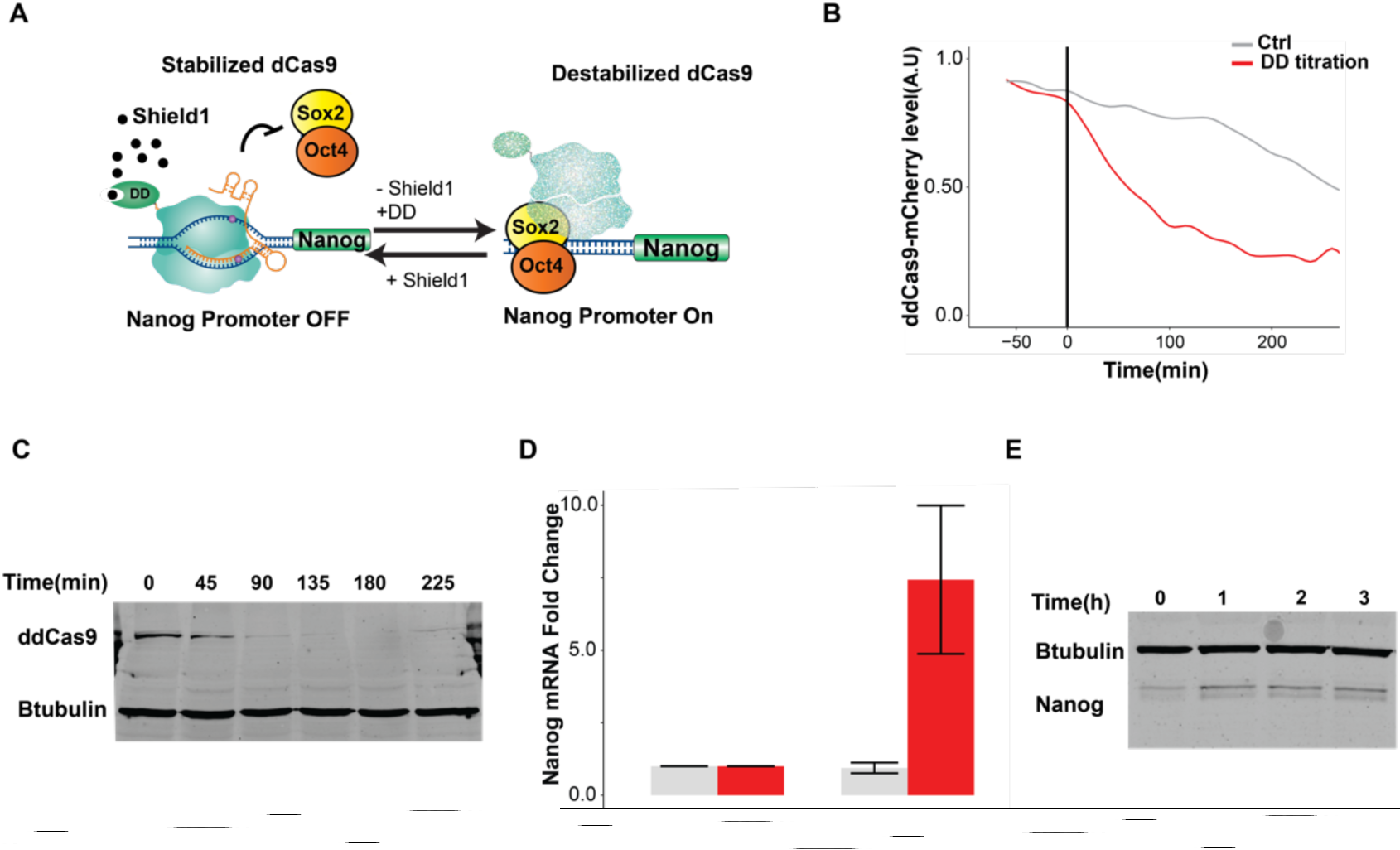
Reversible disruption of TF-DNA interactions with ddCas9. A. Schematic of conditionally-destabilized dCas9 (ddCas9) used for reversible inhibition.
B. Time-course microscopy, and,
C. immunoblot of ddCas9 degradation after Shield1 titration by excess competitive inhibitor protein (DD).
D. Nanog mRNA qRT-PCR, and, E, Immunoblot measurement of Nanog protein after ddCas9 degradation by Shield1 titration.

**Supplementary Figure 1:**
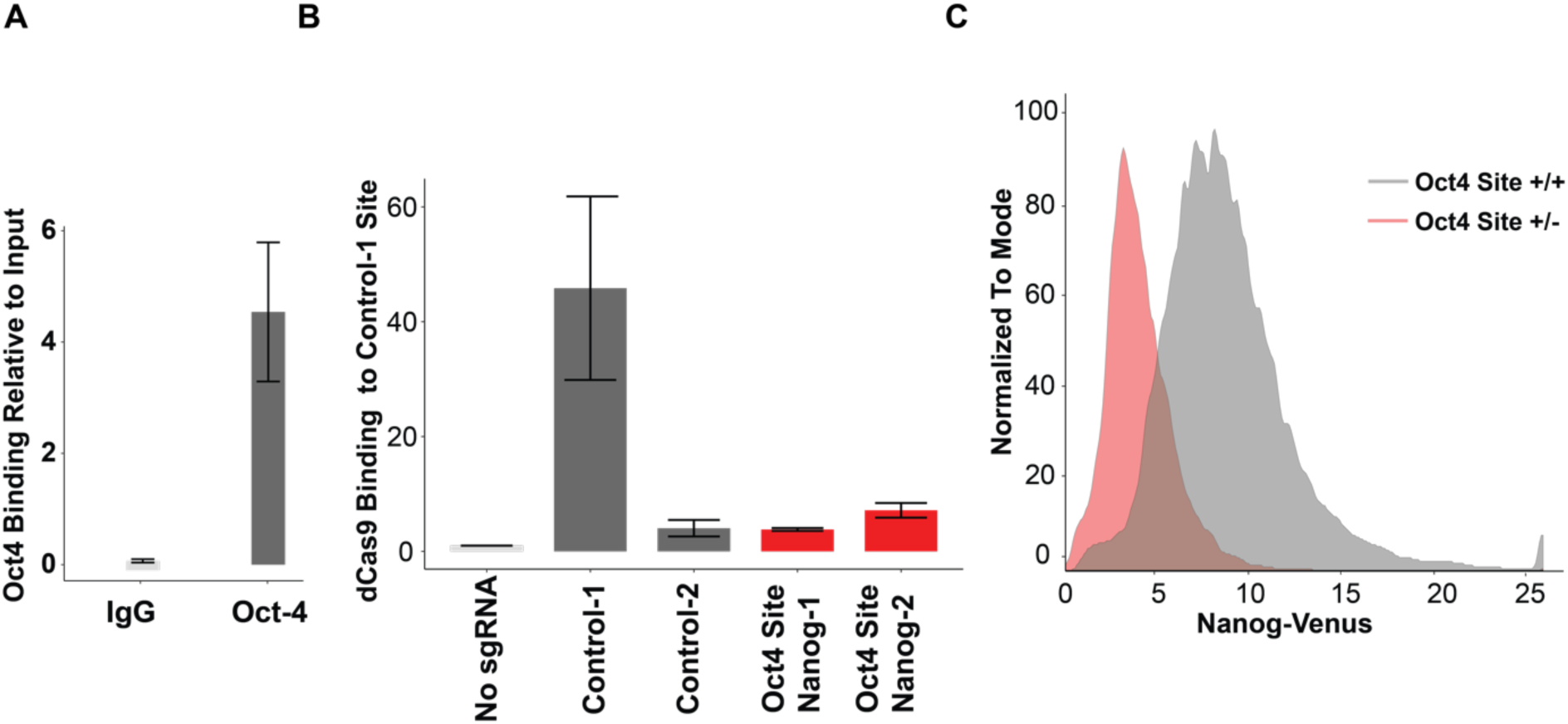
A. Oct4 ChIP-qPCR using primers adjacent to the Oct4 binding site upstream of the *Nanog* gene following pull-down with an Oct4 antibody or IgG control antibody.
B. dCas9 ChIP-qPCR using primers next to the binding site of the control-1 sgRNA (see Fig. 1E).
C. Flow cytometry measurement of Nanog-Venus in wild type (Oct4 site +/+) and heterozygous Oct4 site deletion (Oct4 site +/−) upstream of Nanog-Venus.

**Supplementary Figure 2:**
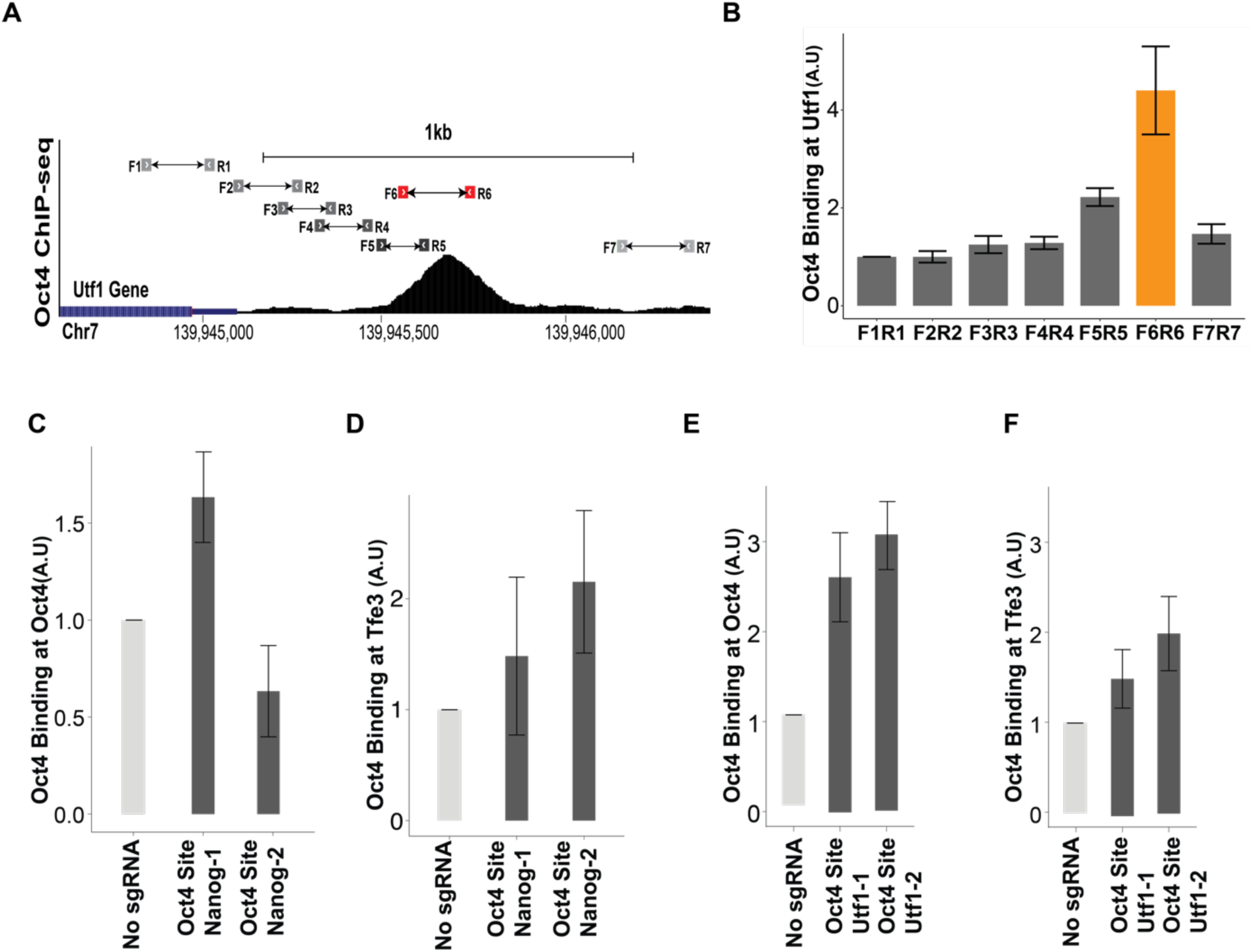
A. Position of ChIP-qPCR primers used to identify the Oct4 binding site in the regulatory sequences downstream of the *Utf1* gene.
B. Oct4 ChIP-qPCR data using a set of overlapping primers to confirm the location of the Oct4 binding site downstream of the *Utf1* gene.
C. ChIP-qPCR analysis of Oct4 binding at an Oct4 site near the *Oct4* gene, or,
D. an Oct4 site near the *Tfe3* gene in ESCs expressing the Oct4-Site Nanog-1 sgRNA or the Oct4-Site Nanog-2 sgRNA.
E. ChIP-qPCR analysis of Oct4 binding at an Oct4 site near the *Oct4* gene, or, D, an Oct4 site near the *Tfe3* gene in ESCs expressing the Oct4-Site Utf1 -1 sgRNA or the Oct4-Site Utf1 -2 sgRNA.

**Supplementary Figure 3:**
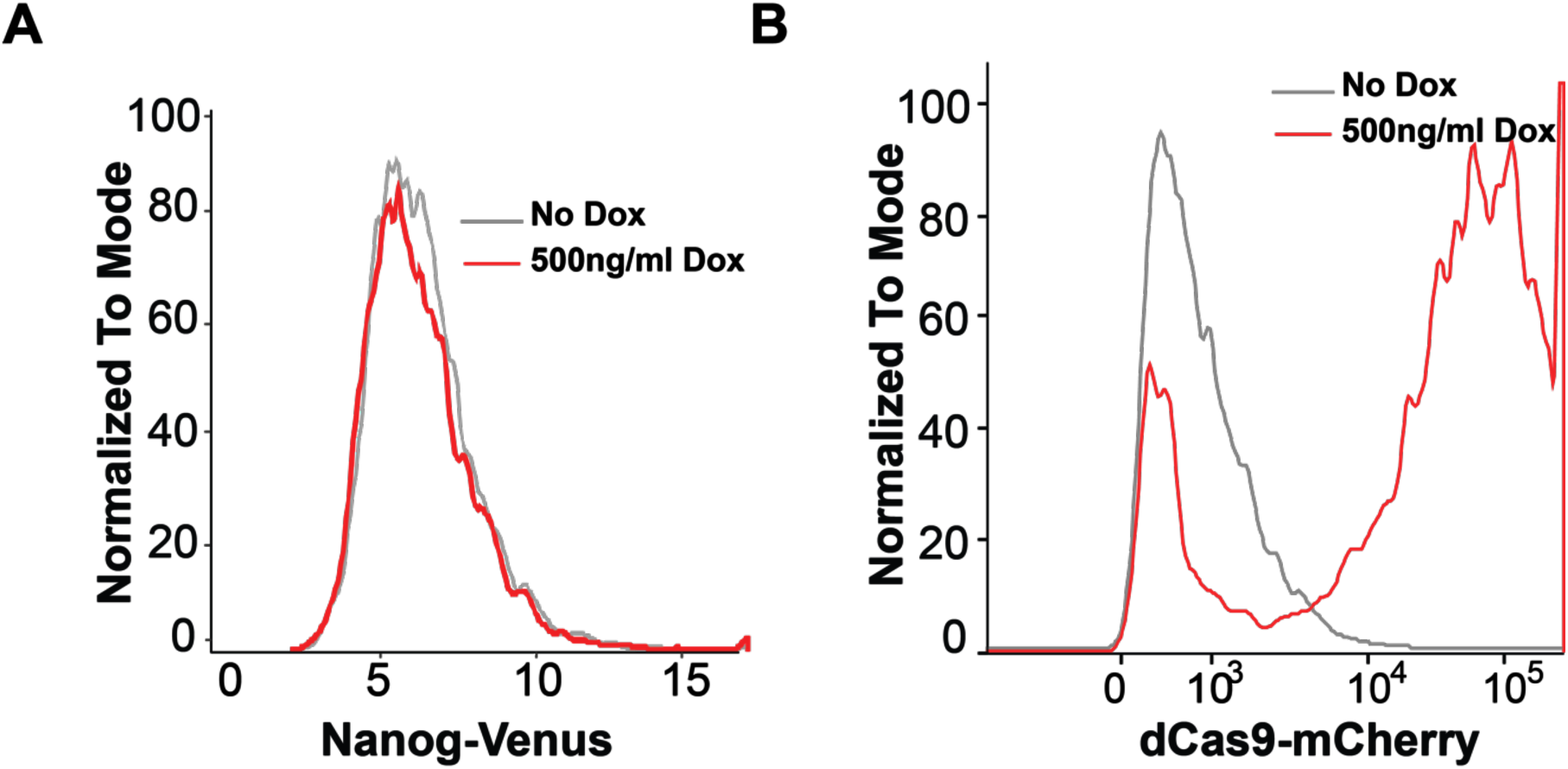
Flow cytometry analysis of Nanog-Venus, and, dCas9-mCherry expression in the Nanog-Venus/dCas9-mCherry ESC line.

**Supplementary Video 1, 2:** Phase contrast time lapse microscopy of the dCas9/Nanog-Venus control cell line without (Video 1) and with expression of the Oct4-Site Nanog-1 sgRNA (Video 2).

**Supplementary Video 3:** Time lapse phase contrast and fluorescent microscopy of degradation of ddCas9-mCherry after Shield1 titration by excess DD protein as a competitive inhibitor (Video 3).

